# Seasonal Changes in Bird Assemblages of a Forest-Steppe Ecotone in North Patagonia

**DOI:** 10.1101/338772

**Authors:** Víctor R. Cueto, Cristian A. Gorosito

## Abstract

We evaluate seasonal variations at the community level, analyzing changes in species richness, species composition and total abundance, and at the species level, evaluating differences in breeding and molting seasonality among bird species in a forest-steppe ecotone of north Patagonia. The bird assemblage showed a low seasonal variation in richness and total abundance, but a great change in species composition between spring-summer and fall-winter. The change in species composition promoted few seasonal variations in richness and total abundance, because they were compensated by the presence of abundant species that visit the area in different seasons. At the species level, resident birds and short distance migrants tended to begin breeding earlier than long distance migrants, and birds began to molt body and flight feathers after breeding. Therefore, we found a low overlap of these two demanding activities. Our results highlight the importance of bird movements in response to seasonal variations in the availability of resources, which promote migration or local displacements of birds.

**Resumen. Cambios estacionales en los ensambles de aves en un ecotono bosque-estepa del norte de Patagonia:** Evaluamos las variaciones estacionales a nivel comunitario, analizando la riqueza de especies, la composición de especies y la abundancia total, y a nivel de las especies, evaluando las diferencias en la estacionalidad reproductiva y de muda entre las especies de aves en un ecotono bosque-estepa del norte de Patagonia. El ensamble de aves mostró pocas variaciones estacionales en la riqueza y abundancia total, pero un notable cambio en la composición de especies entre la primavera-verano y el otoño-invierno. El cambio en la composición de especies promovió pocas variaciones estaciones en la riqueza y la abundancia total, porque fueron compensadas por la abundancia de las especies que visitan el área en las diferentes estaciones. A nivel de las especies, las aves residentes y migrantes de corta distancia tendieron a comenzar la reproducción antes que las migrantes de larga distancia, y las aves comenzaron la muda de plumas del cuerpo y del ala al finalizar la reproducción. Por lo cual encontramos una baja superposición de estas dos demandantes actividades. Nuestros resultados remarcan la importancia del movimiento de las aves en respuesta a las variaciones en la disponibilidad de recursos, que promueven la migración o el desplazamiento local de las aves.

## INTRODUCTION

Seasonal changes in the richness and abundance of bird species assemblages are characteristics of temperate forests (Wiens 1989), in part because some species migrate to avoid harsh climate conditions and reduced food supply during winter months (Newton 2008). In temperate regions of South America several studies have been conducted on the seasonality of bird assemblages (e.g., Capurro & Bucher 1986, Marone 1992, Rozzi et al. 1996, Cueto & Lopez de Casenave 2000, Becerra & Grigera 2005, Ippi et al. 2009, Kelt et al. 2012), showing different patterns of seasonal changes, independent of geographic location. In the Chaco Forest (Capurro & Bucher 1986), Monte Desert (Marone 1992), Chilean Mediterranean Scrublands (Kelt et al. 2012) and northern Patagonian Forest (Rozzi et al. 1996, Becerra & Grigera 2005), richness values were lower during autumn-winter than in spring-summer, because of the migration of several species. Conversely, in Chilean southernmost Patagonian forest (Ippi et al. 2009) and in the Argentinean Coastal Woodlands (Cueto & Lopez de Casenave 2000) there was little variation in richness, because bird assemblages were composed by few migrant species. Although bird migration in South America is not considered to be as common as it is in the North Hemisphere (Elphick 2007), those studies on avian seasonality have shown the importance of migratory species in the structure of avian assemblages across different South American ecosystems.

General aspects about the origin and biogeography of birds inhabiting Patagonian forests are well known (Vuilleumier 1985). Also, there was an increase in the number of studies on bird communities in different forest types (Rozzi et al. 1996, Becerra & Grigera 2005, Grigera & Pavic 2007, Lantschner et al. 2008, Ippi et al. 2009). However, life history aspects of bird species have been less studied. For example, despite previous studies on patterns of feather molt in the tropics of South America (e.g., Marini & Durães 2001, Ryder & Wolfe 2009, Silveira & Marini 2012, Jahn et al. 2016), almost nothing is known about this aspect of bird species in the Patagonian forest. Here, we evaluate seasonal variations in richness, species composition and total abundance at community level, and differences in breeding and molting seasonality among bird species at species level in a forest-steppe ecotone of North Patagonia. Considering the climate seasonality in Patagonia, with harsher weather conditions during fall-winter than spring-summer, we expect lower richness and total abundance and no breeding and molting activity during autumn and winter than in spring and summer seasons.

## METHODS

### Study site

We conducted our study at Cañadón Florido Ranch (42° 55’ S, 71° 21’ W), Chubut Province, Argentina. The vegetation of the area corresponds to the Valdivian Forest Province of the Andean Region (Morrone 2001). We studied bird assemblages in a forest dominated by *Maytenus boaria* trees, commonly called Maitenales (Dimitri 1972). This forest is common in valleys and slope of hills in the eastern portion of Patagonian forest in Chubut Province, and forms part of the ecotone between the forest and the steppe (Kitzberger 2012). The forest canopy is low, averaging a height of 5 m. The understory is mainly dominated by *Berberis microphylla* shrubs. The climate is characterized by cold and wet winters and mild but dry summers. Most precipitation falls as rain and snow during fall and winter (April - September). Annual mean precipitation in the region is 484 mm. Summer and winter mean temperatures are 17° and 2.5 °C, respectively.

### Bird survey

We caught birds with mist nets within a 20-ha plot (Fig. 1) from October 2015 to March 2017. Nets were 12 m long with a 38 mm mesh size and were opened for 4–5 h after sunrise. We set two groups of 10 nets within the plot. Each group was set in different forest patches, 200 m apart (Fig. 1). Nets within each group were set 50 to 70 m apart and opened at least once per month. The order of net group sampling was set randomly, with 10 or 15 days between sampling events, depending on weather conditions (i.e., avoiding rainy or windy days). Due to adverse weather conditions, we sampled birds only once during June 2016 and February 2017, and because of logistic problems, we did not sample birds during April and July 2016. Our sampling effort consisted of 49 net-days and 2469 net-hours. In Table 1 we indicate the number of sampling days and the net hours of sampling on each month. We banded all birds with aluminum leg bands and took five morphological measurements: body mass, wing length, tail length, tarsus length, and bill length (from the anterior end of the nostril to the bill tip). We used a digital scale (± 0.1 g) to record body mass, a wing ruler (± 1 mm) for wing and tail measurements, and a digital caliper (± 0.01 mm) for tarsus and bill measurements. We scored body molt intensity on a 5-point scale, from none to heavy (none = no feathers molting; trace = few feathers molting; light = involving more than one feather tract; medium = half of body feathers molting; heavy = most/all body feathers molting; Ralph et al. 1993). We scored remige molt by noting which primary feather and secondary feather was molting on each wing (Ralph et al. 1993). When classifying flight feather molt, we excluded birds with adventitious molt, following Wolfe et al. (2010), who considered symmetrical molt of the first primary feather to indicate the start of a molt cycle. Thus, we classified birds as being in remige molt when they were molting at least one primary feather on each wing. We scored the level of brood patch development on a 5-point scale (0 = no patch; 1 = smooth; 2 = vascularized; 3 = wrinkled; 4 = molting; Ralph et al. 1993). We scored development of the cloacal protuberance on a 4-point scale (0 = none; 1 = small, 2 = medium; 3 = large; Ralph et al. 1993). We determined sex of each individual based on characteristics of the plumage (Canevari et al. 1992), morphological differences (e.g., Cueto et al. 2015, Pyle et al. 2015), or the presence of a brood patch for females and a cloacal protuberance for males of species with sexually monomorphic plumage.

**Table 1.**
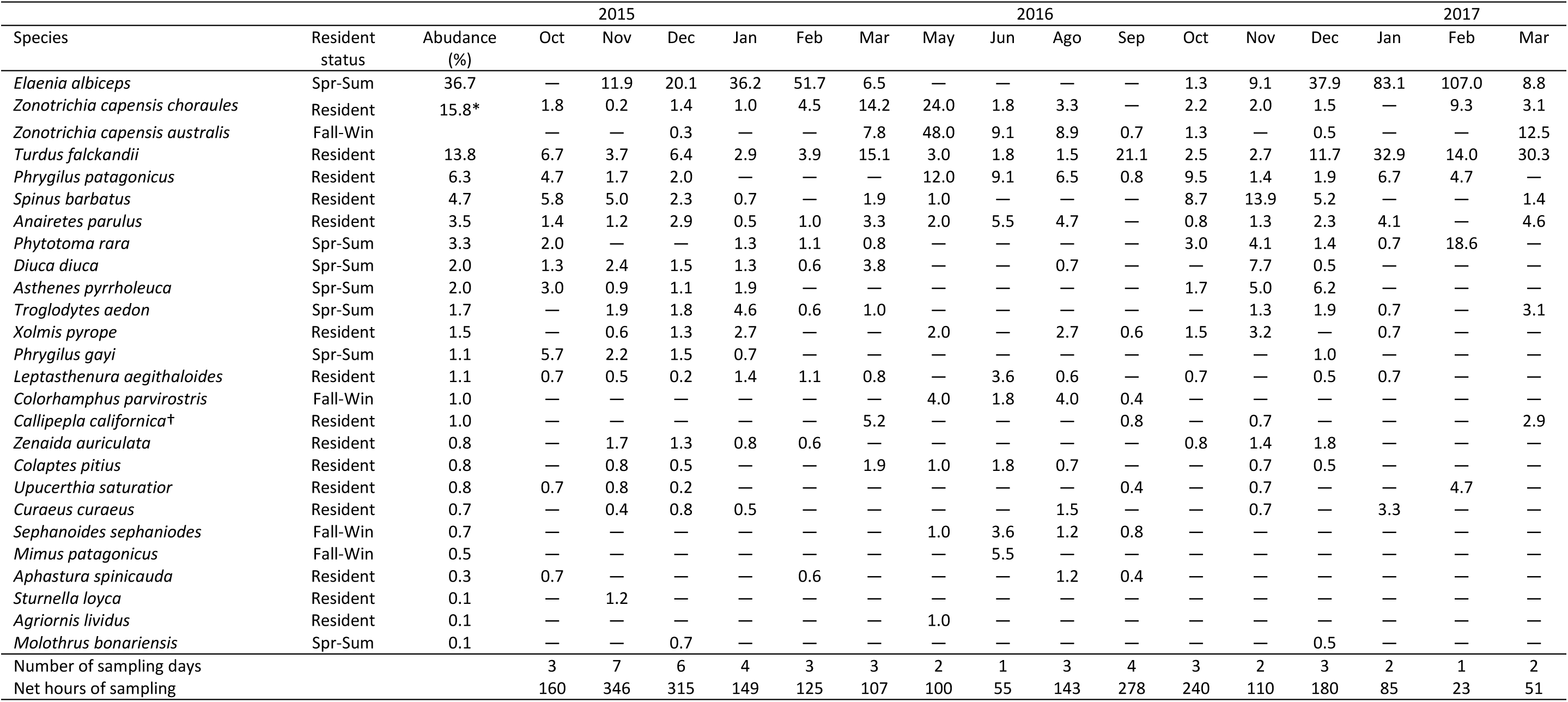
Monthly abundance (individuals captured per_100 net hours) for each bird species captured during the 2015-2017 period in a forest-steppe ecotone at Cañadón Florido Ranch, Chubut Province, Argentina. For ‘Resident status’, ‘Spr-Sum’ indicates species that visit Cañadón Florido in spring-summer and Fall-Win during fall-winter months. Species are listed in order of their percent contribution to average total abundance during the study period. The symbol “—” denotes zero individuals captured. * percentage corresponds to both subspecies of *Zonotrichia capensis* in each month. † Introduced species from North America.

**Figure 1.**
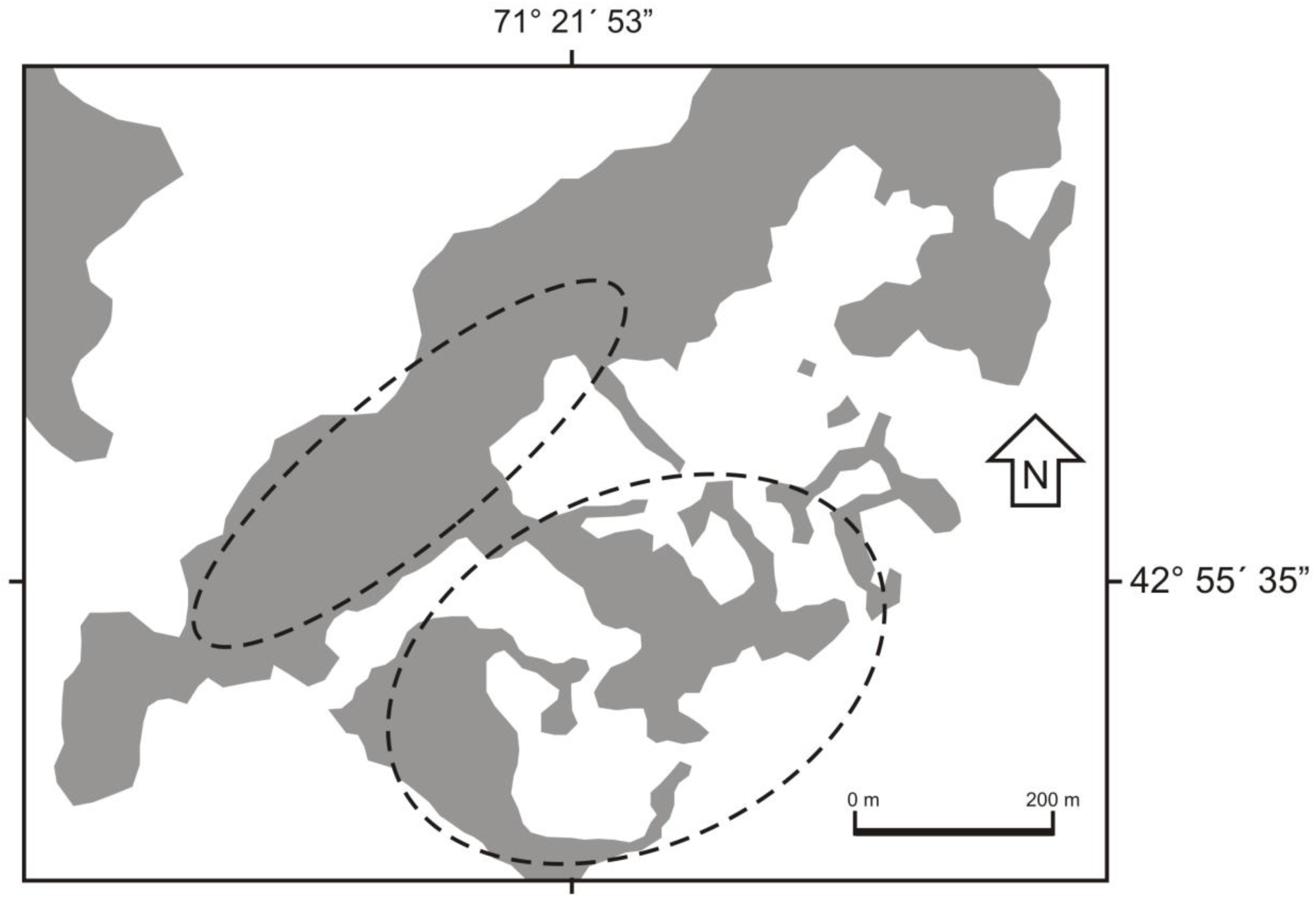
Study site map showing the forest areas (in grey) at Cañadón Florido Ranch, Chubut Province, Argentina. The two ellipses indicate the zone where mist nets were installed in the area.

### Climate patterns

To assess the potential effect of climate on bird assemblage seasonality, we analyzed monthly variations on temperature and precipitation at the nearest weather station to our study site. Data were available from the weather station “RÍo Percy” (42° 51’S, 71° 25’ W, belonging to Hidroeléctrica Futaleufú), located at 9 km from Cañadón Florido Ranch. We considered only mean temperature, because maximum mean temperature and minimum mean temperature were strongly correlated with this variable (r = 0.97 and r = 0.96, respectively).

### Data analysis

Abundance of each species was estimated using capture rate as the number of captures per 100 net-hours (Karr 1981). Although mist net sampling could underrepresent species richness because of a lower probability to capture canopy (Mallory et al. 2004), we considered that our estimation of richness is unbiased. First, the canopy height of the woodlands in our study area is around 5 m and our nets have a height of 3 m. Second, we have been working in the study area since November 2013 (see Bravo et al. 2017), and every time we visited the area, we recorded all bird species observed or heard. With the exception of raptors (e.g., *Milvago chimango, Polyborus plancus, Bubo magellanicus*) and bird species common in open habitats (e.g., *Hymenops perspicillatus, Lessonia rufa, Muscisaxicola macloviana*), we captured all species that we had recorded in the forest of our study area. Third, the species lists we recorded are similar to others reported in studies of the eastern zone of north Patagonian forests (e.g., Ralph 1985, Grigera & Pavic 2007).

Assemblage structure was evaluated through seasonal changes in total abundance, species richness and species composition. Total abundance was the sum of all new individuals captured each month (i.e., we did not include recaptured individuals during each month, but were considered when recaptured between months). Richness was estimated as the number of species captured each month. Species composition variations among months was evaluated through Cluster Analysis (with the statistical program InfoStat 2009). Months were classified based on the abundance of the species, using the Bray-Curtis dissimilarity index, and the dendrogram was constructed with the UPGMA algorithm (Sneath & Sokal 1973). We used the Cophenetic Correlation Coefficient to measure the agreement between the dendrogram and the original dissimilarity matrix (Sneath & Sokal, 1973). To determine the level of dissimilarity defining groups in the dendrogram, we considered the criteria of mean dissimilarity among all month pairs (i.e., the average of all values in the original dissimilarity matrix; e.g., Holmes, 1981). In our Cluster Analysis we obtain a high Cophenetic Correlation Coefficients (r = 0.79) indicating that the dendrogram is an accurate representation of the structure of the original dissimilarity matrices. Seasonality in richness and total abundance were evaluated through correlation analysis with monthly variations in temperature and precipitation, using Pearson Correlation tests.

Actively breeding birds were identified by presence of a smooth or vascularized brood patch in females or a small, medium or large cloacal protuberance in males. At the assemblage level, we estimated the average percentage of females and males of all species actively breeding in each month, and the average percentage of individuals of all species molting body and wing feathers in each month. We estimated the average abundance of females and males of each species actively breeding in each month, and the average abundance of individuals of each species molting body and wing feathers in each month.

## RESULTS

We captured 25 species and a total of 969 individuals were banded (Table 1). Little seasonal variation was evident in richness and total abundance, although climate showed a seasonal pattern, with lower temperatures during fall and winter and higher precipitations in winter (Fig. 2). Richness was higher in spring (October, November and December), but during other seasons show few changes (Fig. 2b). Total abundance did not show seasonality (Fig. 2c), the lowest value was recorded in September, and during January and February of the summer 2017 there was an abrupt increase, with more than twice the abundance recorded during the same months of the summer 2016 (Fig. 2c). Richness was not correlated with monthly variations in temperature and precipitation (r = 0.21, P = 0.43, n =16 and r = -0.29, P = 0.27, n= 16, respectively). Total abundance also was not correlated with temperature and precipitation variations (r = 0.40, P = 0.13, n = 16 and r = 0.01, P = 0.97, n = 16, respectively). In contrast, species composition shows seasonality. Mean dissimilarity among months based on species abundance distinguished three groups in the dendrogram (Fig. 3). One group was constituted by spring-summer months, the second group by winter months, and a third group formed by March and September, i.e., transitional months from summer to autumn (March) and from winter to spring (September). Spring-summer group was mainly composed by *Elaenia albiceps, Phytotoma rara, Diuca diuca, Asthenes pyrrholeuca* and *Troglodytes aedon*, and winter group by *Zonotrichia capensis, Colorhamphus parvirostris* and *Sephanoides sephaniodes* (Table 1). The transitional group was mainly characterized for the abundance of *Turdus falcklandii* (Table 1).

**Figure 2.**
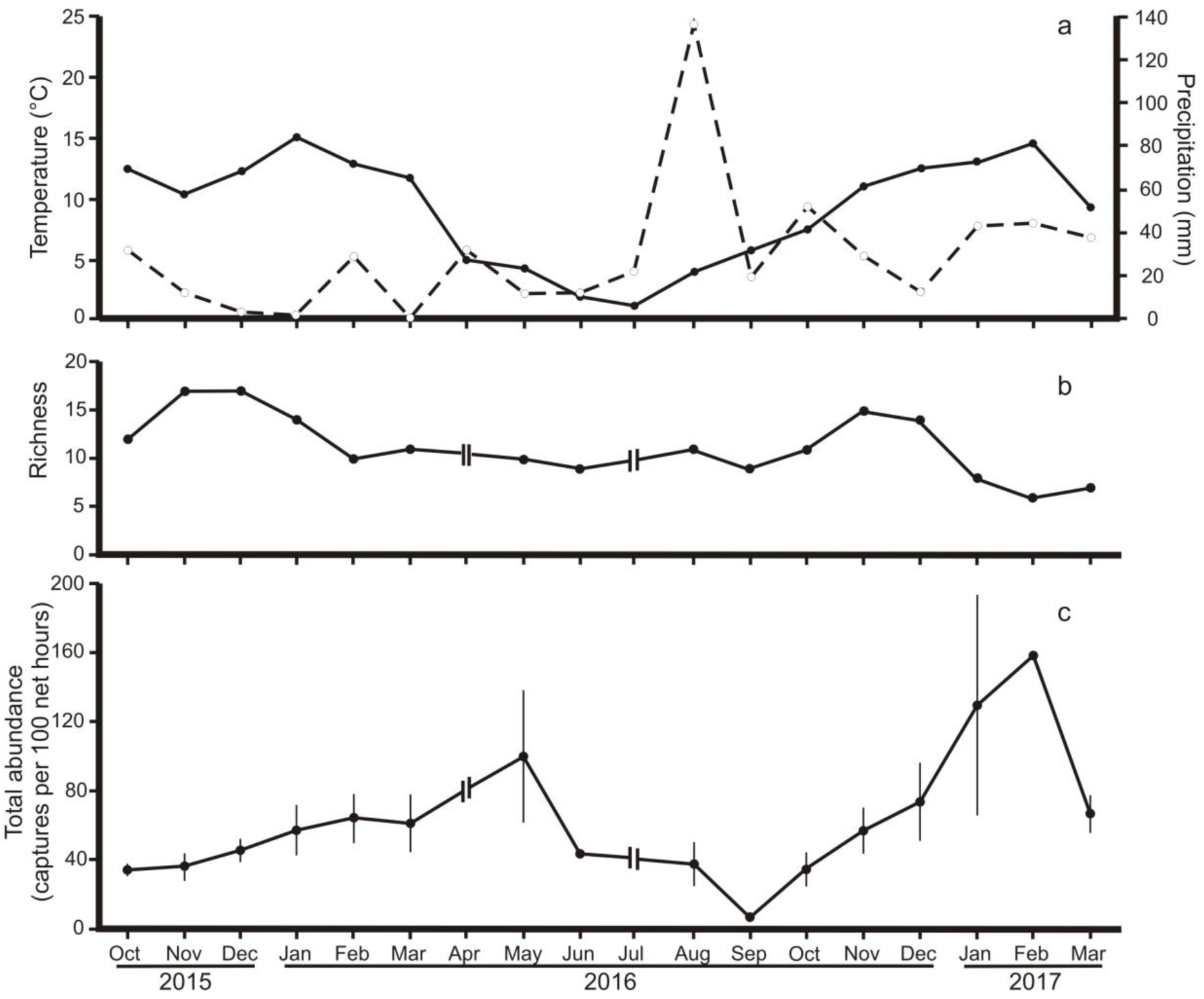
Monthly changes in mean temperature (solid line) and precipitation (dashed line) (a), richness (b) and total abundance (individuals captured per 100 net hours) (c) in a forest-steppe ecotone at Cañadón Florido Ranch, Chubut Province, Argentina.

**Figure 3.**
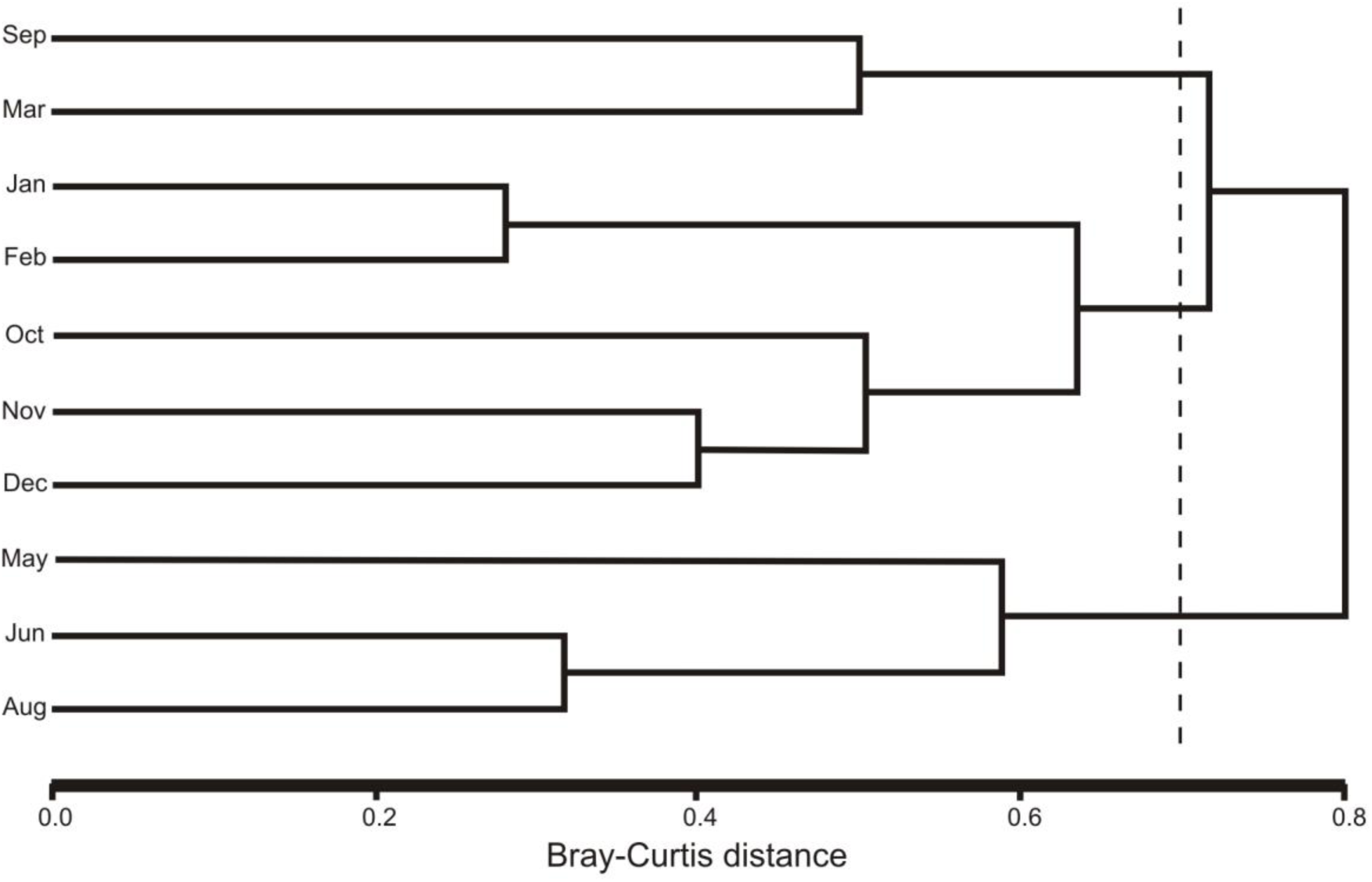
Cluster Analysis based on bird species abundance from a forest-steppe ecotone (Cañadón Florido Ranch, Chubut Province, Argentina), showing month groups. Dashed line indicates mean dissimilarity among all month pairs.

More than 35 % of all captured individuals were *Elaenia albiceps* (Table 1). This species began to be captured during the last days of October, with captures through March (Table 1). During the second summer we had a high capture rate of *Elaenia albiceps* (January and February 2017, Table 1). The second most abundant species was *Zonotrichia capensis* with more than 15% of all captured individuals (Table 1). We captured two subspecies: *Zonotrichia capensis choraules* and *Zonotrichia capensis australis*. The former was present all year-round, although more abundant during fall (mainly in May, Table 1), while the latter was abundant during fall and winter months (Table 1). The other two more abundant species were *Turdus falcklandii* and *Phrygilus patagonicus* (Table 1), and both were present all year-round. In summary, these four species represent more than 70% of the annual average total abundance in the bird assemblage.

There were another five species present only during spring-summer and three during fall-winter. Spring-summer visitors were *Phytotoma rara, Diuca diuca, Asthenes pyrrholeuca, Troglodytes aedon* and *Phrygilus gayi*, and fall-winter visitors were *Colorhamphus parvirostris, Sephanoides sephaniodes* and *Mimus patagonicus* (Table 1). The rest of the species were present all year-round or had no clear seasonal pattern. For example, *Spinus barbatus* was very abundant only during spring, and was scarce in summer and fall and absent during winter (Table 1).

The breeding season lasted from September to March (Table 2). Males were the first individuals captured in breeding condition. The extreme case was a *Turdus falckandii* male captured in September (Table 2). Males of the rest of the species in breeding condition began to be captured in October (Table 2), with the exception of *Elaenia albiceps, Diuca diuca* and *Troglodytes aedon* males, whose first captures began in November (Table 2). Females in breeding condition were captured beginning in November, but *Elaenia albiceps* and *Diuca diuca* females in breeding condition were first captured in December and January, respectively (Table 2).

**Table 2.**
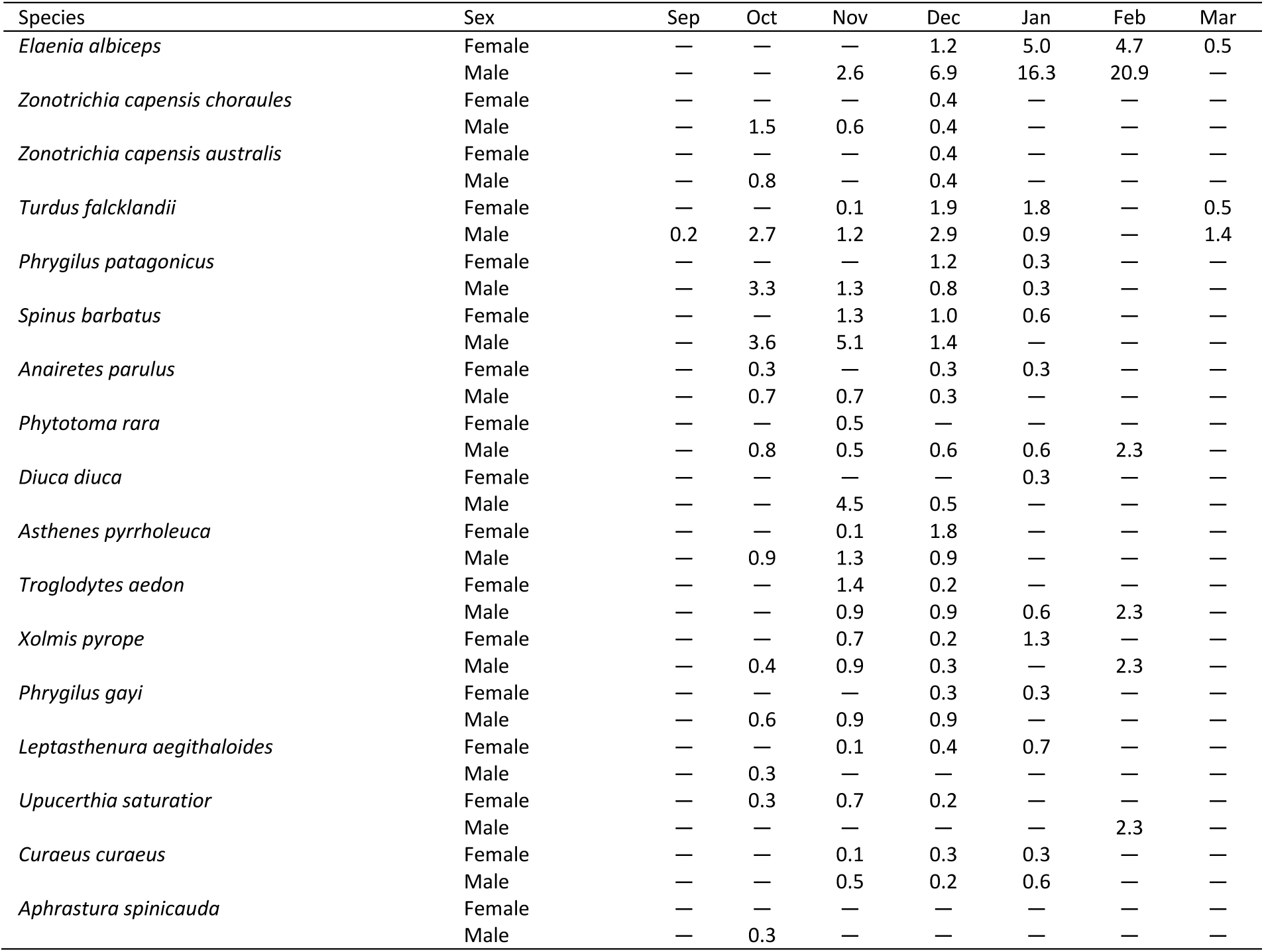
Average monthly abundance (individuals captured per 100 net-hours) of females and males in breeding condition across bird species in a forest-steppe ecotone at Cañadón Florido Ranch, Chubut Province, Argentina. Species are listed following the order in Table 1. The symbol “—” denotes zero individuals captured in breeding conditions.

Even though almost all species began molting body and wing feathers during summer months (mainly in February and March, Table 3), we recorded *Zenaida auriculata* and *Xolmis pyrope* molting feathers in November and December, respectively (Table 3). Two migrant species, *Troglodytes aedon* and *Diuca diuca* molted body and wing feathers (Table 3), but we did not capture any *Elaenia albiceps* molting. There was little overlap between breeding and molting events (Fig. 4). The percentage of breeding females was higher in December and January, while there was a similar percentage of breeding males from October to February (Fig. 4). In contrast, the highest percentage of individuals molting body and wing feathers was in March (Fig. 4).

**Table 3.**
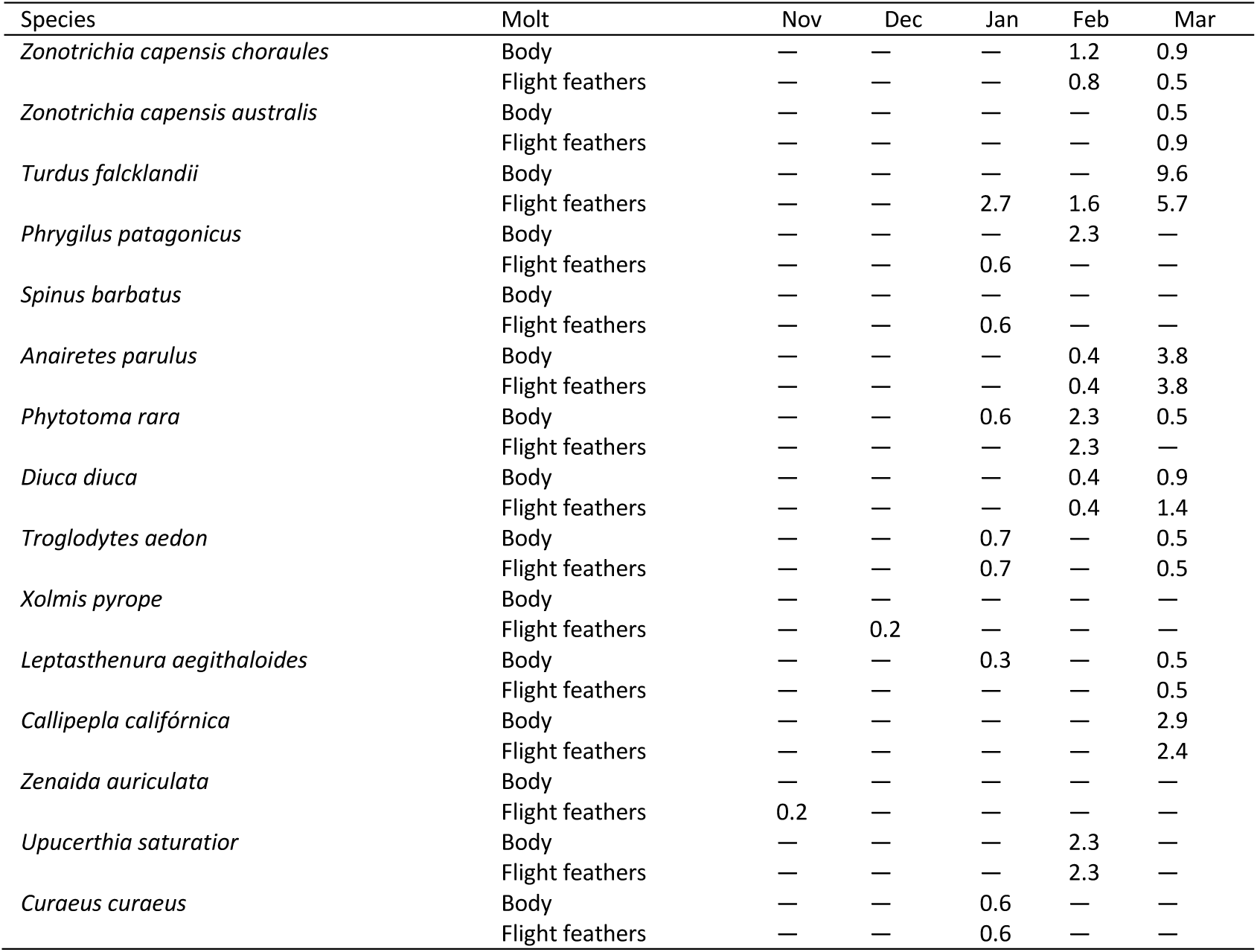
Average monthly abundance of molting individuals (individuals captured per 100 net-hours) of each bird species in a forest-steppe ecotone at Cañadón Florido Ranch, Chubut Province, Argentina. Species are listed following the order in Table 1. The symbol “—” denotes zero individuals captured in molting conditions.

| Species | Molt | Nov | Dec | Jan | Feb | Mar |
| --- | --- | --- | --- | --- | --- | --- |
| *Zonotrichia capensis choraules* | Body | — | — | — | 1.2 | 0.9 |
|  | Flight feathers | — | — | — | 0.8 | 0.5 |
| *Zonotrichia capensis australis* | Body | — | — | — | — | 0.5 |
|  | Flight feathers | — | — | — | — | 0.9 |
| *Turdus falcklandii* | Body | — | — | — | — | 9.6 |
|  | Flight feathers | — | — | 2.7 | 1.6 | 5.7 |
| *Phrygilus patagonicus* | Body | — | — | — | 2.3 | — |
|  | Flight feathers | — | — | 0.6 | — | — |
| *Spinus barbatus* | Body | — | — | — | — | — |
|  | Flight feathers | — | — | 0.6 | — | — |
| *Anairetes parulus* | Body | — | — | — | 0.4 | 3.8 |
|  | Flight feathers | — | — | — | 0.4 | 3.8 |
| *Phytotoma rara* | Body | — | — | 0.6 | 2.3 | 0.5 |
|  | Flight feathers | — | — | — | 2.3 | — |
| *Diuca diuca* | Body | — | — | — | 0.4 | 0.9 |
|  | Flight feathers | — | — | — | 0.4 | 1.4 |
| *Troglodytes aedon* | Body | — | — | 0.7 | — | 0.5 |
|  | Flight feathers | — | — | 0.7 | — | 0.5 |
| *Xolmis pyrope* | Body | — | — | — | — | — |
|  | Flight feathers | — | 0.2 | — | — | — |
| *Leptasthenura aegithaloides* | Body | — | — | 0.3 | — | 0.5 |
|  | Flight feathers | — | — | — | — | 0.5 |
| *Callipepla californica* | Body | — | — | — | — | 2.9 |
|  | Flight feathers | — | — | — | — | 2.4 |
| *Zenaida auriculata* | Body | — | — | — | — | — |
|  | Flight feathers | 0.2 | — | — | — | — |
| *Upucerthia saturator* | Body | — | — | — | 2.3 | — |
|  | Flight feathers | — | — | — | 2.3 | — |
| *Curaeus curaeus* | Body | — | — | 0.6 | — | — |
|  | Flight feathers | — | — | 0.6 | — | — |

**Figure 4.**
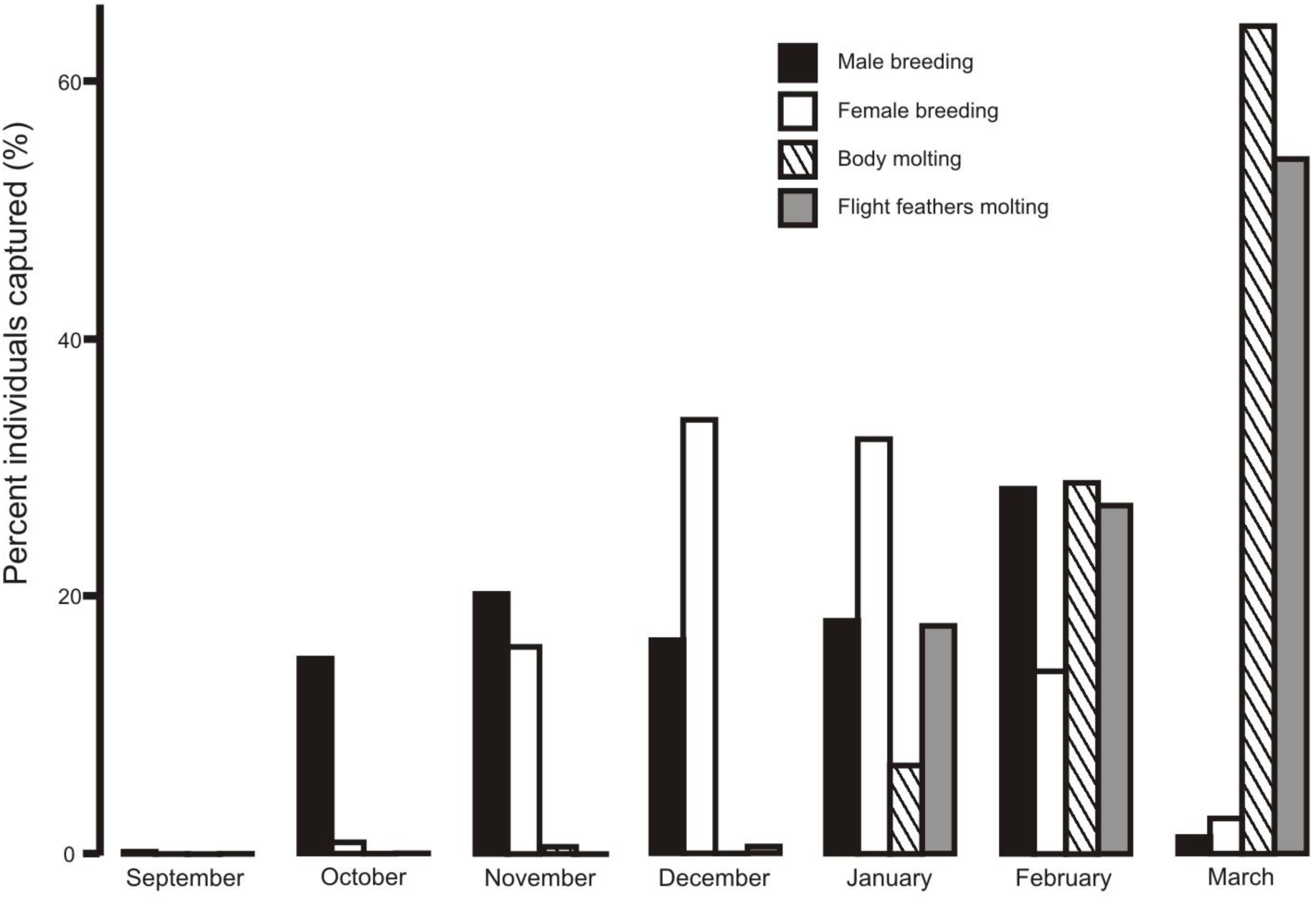
Monthly changes in the percentage of individuals in breeding and molting condition in a forest-steppe ecotone at Cañadón Florido Ranch, Chubut Province, Argentina.

## DISCUSSION

The bird assemblage at Cañadón Florido did not show a seasonal pattern in richness and total abundance, contrary to our expectation given the strong climatic seasonality in this temperate zone of South America. Our results contrast with seasonal patterns of community parameters in temperate forests of Patagonia at the same latitude of our study site (i.e., lower richness and bird abundance during autumn-winter than in spring-summer, Rozzi et al. 1996), but are similar to that reported for the most southern forest of South America (Ippi et al. 2009). These contrasting results could be related to differences in location, forest composition, climatic conditions and bird assemblage composition. For example, in the southern forests of Chile one third of the bird assemblage was composed by migratory species (Ippi et al. 2009), but about a half of the species in forests further north in Chile were found to be migrants (Jaksic & Feinsinger 1991). In our study site, a forest-steppe ecotone of North Patagonia, 42 % of the species are migrants, but there are seven species that visit the zone during spring-summer and four that are present in fall-winter. This seasonal turnover of species that visit Cañadón Florido results in low seasonal variation of richness and total abundance. Also, the species turnover of bird assemblage in Cañadón Florido could be the reason of the lack of association among precipitation and temperature with bird richness and total abundance.

Visitors during fall-winter could be a result of the displacement of several species from mountain forests or from the steppe. For example, Rozzi et al. (1996) suggested that *Sephanoides sephaniodes* moves to lowland forest in winter, and in our study area we captured *Mimus patagonicus*, which is common in the steppe (Llanos et al. 2017). Moreover, the presence of *Colorhamphus parvirostris* during fall-winter could be related to movements from highland forest, similar to the displacement patterns of *Sephanoides sephaniodes*. *Colorhamphus parvirostris* is a partial migrant that has a year-round presence in the northern portion of its breeding distribution (Chesser & Marin 1994). Another partial migrant that visits our study site during fall-winter is *Zonotrichia capensis australis*. A similar pattern was recorded by Keve & Kovács (1971) for this subspecies in localities near El Bolsón and Bariloche (Argentina).

*Zonotrichica capensis australis* breeds mainly in southern Argentina and Chile (King 1974, Piloni 2002, Ortiz & Capllonch 2011), and moves to northern latitudes in winter (Ortiz & Capllonch 2011, Sagario et al. 2014). During May we had a high capture rate of this subspecies, but during the following fall and winter months its capture rate was low. Thus, these birds could be on passage migration to lower latitudes in May. During spring and summer, we captured very few individuals of this subspecies in breeding condition, so at least a small population of this subspecies potentially breeds at our study site. During spring there is strong change in species composition due to the departure of winter visitors and then arrival of long-and short-distance migrants, as *Elaenia albiceps* (a long-distance migrant, Bravo et al. 2017), *Phytotoma rutile, Diuca diuca, Asthenes pyrrholeuca* and *Troglodytes aedon* (short-distance migrants).

Total bird abundance did not show strong seasonal changes because it was compensated by spring-summer and fall-winter visitors. During spring and summer, *Elaenia albiceps* was the most abundant species in the Maitenal forest at Cañadón Florido ranch. This species arrived during the last weeks of October and the last individuals were captured in late March. Its high abundance during those months is also common in the eastern and western forests of Patagonia (e.g., Ralph 1985, Grigera et al. 1994, Rozzi et al. 1996, Jiménez 2000, Ippi et al. 2009). During fall and winter two subspecies of *Zonotrichia capensis* were the most abundant, and their high capture rates compensated the reduction in abundance due to departure of spring-summer visitors. *Zonotrichia capensis choraules* was resident all year-round, but its high capture rate was between March and August, and during the same months the partial migrant *Zonotrichia capensis australis* was also very abundant. *Zonotrichia capensis choraules* is resident in other localities of northern Patagonia (Keve & Kovács 1971), and both subspecies form flocks of hundreds of individuals during fall-winter in northern Patagonia (Keve & Kovács 1971) and in the region of the Monte desert (Sagario et al. 2014, Zarco & Cueto 2017). During the second summer of our study (mainly January and February 2017), total abundance increased notably. This increase was due to a high capture rate of *Elaenia albiceps* and *Turdus falkclandii* during that summer. Fruit abundance of *Berberis microphylla* and *Maytenus boaria* was higher in the summer 2017 than 2016 (Gorosito C.A. & Cueto V.R., unpublished data). As these two species consume large quantities of fruit from those plant species (Amico & Aisen 2005), the high abundance of fruits could be the reason for their high capture rate during the summer 2017. Abundance of *Molothrus bonariensis* was low at Cañadón Florido ranch. This species is a generalist brood parasite that exerts significant impacts on the breeding success of its hosts (Reboreda et al. 2003). We recorded only five nests parasitized by *Molothrus bonariensis*, out of the 145 nests we found of 12 species (Gorosito C.A., unpublished data). Notably, all parasitized nests were of *Diuca diuca*. Therefore, at our study site, the effect of brood parasitism on breeding success of birds could be lower than that reported for northern localities (Reboreda et al. 2003).

Resident birds and short distance migrants tend to begin breeding earlier than long distance migrants (Murphy 1989). In Cañadón Florido we found the same pattern, since *Turdus falcklandii* males were in breeding condition as early as September, and males of the other resident and several short distance migrant species were in breeding condition in October. In contrast, captures of males in breeding condition of the long distance migrant *Elaenia albiceps* began in November. Females showed the same pattern than males, but differed over time. Females of resident and almost all short distance migrant species were in breeding condition in November, but captures of *Elaenia albiceps* females in breeding condition began in December. Similar differences in the beginning of reproduction between resident and long-distance migrants were found in species breeding in Monte desert (Mezquida 2002).

At Cañadón Florido, birds began to molt body and flight feathers in the last months of summer, with the exception of *Elaenia albiceps*, which did not molt. This long-distance migrant appears to molt in its wintering areas (Pyle et al. 2015). For example, birds molting the outer primaries were captured in Chapada dos Guimarães, Mato Grosso, Brazil (Guaraldo, A.C., personal communication). Feather molt is an energetically expensive activity within a bird’s annual cycle (Lindström et al. 1993, Murphy 1996).

Therefore, it is more common that breeding and molting overlap in the tropics (Foster 1974, Johnson et al. 2012, de Araujo et al. 2017), whereas in temperate zones there is a low overlap between these two demanding activities (Jenni & Winkler 1994, Murphy 1996, Jahn et al. 2017). At our study site, we found this last pattern, as birds began to molt after breeding, and during a period of favorable climatic conditions. Hence, at the end of the summer, food availability was presumably sufficient for birds to pay the energetic cost of molting feathers.

In summary, the bird assemblage at Cañadón Florido showed lower seasonality in overall richness and total abundance, because of a turnover in the composition of species between spring-summer and fall-winter. These changes highlight the importance of bird movements in response to seasonal variation in resource availability that promote migration or local displacements of birds.

## ACKNOWLEDGMENTS

We thank the Roberts family for allowing us to work at Cañadón Florido. For constructive reviews of the manuscript, we thank Alex Jahn and two anonymous referees. Research carried out at Cañadón Florido was founded by the National Geographic Society (USA) and the Bergstrom Award of the Association of Field Ornithologists (USA). Birds were captured with permission of Dirección de Fauna y Flora Silvestre, Provincia del Chubut (Argentina).

